# A random-dot kinematogram for web-based vision research

**DOI:** 10.1101/192377

**Authors:** Sivananda Rajananda, Hakwan Lau, Brian Odegaard

## Abstract

Web-­based experiments using visual stimuli have become increasingly common in recent years, but many frequently-used stimuli in vision research have yet to be developed for online platforms. Here, we introduce the first open access random-dot kinematogram (RDK) for use in web browsers. This fully customizable RDK offers options to implement several different types of noise (random position, random walk, random direction) and parameters to control aperture shape, coherence level, the number of dots, and other features. We include links to commented JavaScript code for easy implementation in web-based experiments, as well as an example of how this stimulus can be integrated as a plugin with a JavaScript library for online studies (jsPsych).

## (1) Overview

### Introduction

Over the last several decades, random-dot kinematograms (RDKs) have emerged as an effective psychophysical stimulus to evaluate low-level motion processing [1–5]. While different versions of this type of stimulus have been created across several software platforms (e.g., MATLAB [6] and PsychoPy [7,8]), currently, no implementation of this stimulus exists for public use in web-based experiments where the subjects complete the experimental task using web browsers. Recently, our research group developed an RDK in JavaScript to be used for presentation in standard web browsers. This RDK incorporates several distinct features that have emerged in different versions used by researchers, and allows users to customize noise types and other parameters for different paradigms. In this short article, we explain a few important elements of this stimulus and provide links to both raw code and web-based examples to guide implementation for vision scientists in future experiments.

### Implementation and architecture

In RDKs, a certain percentage of dots are designated as “signal” to move in one coherent direction, and the remaining percentage of dots are designated as “noise” to move in random directions. However, as noted by [9], several options exist regarding how signal and noise can be drawn in frame-by-frame presentations. To create the “signal,” dots can either move in the direction of coherent motion in *all* frames (referred to as the “same” rule), or move in the direction of coherent motion in only a specified proportion of frames (referred to as the “different” rule). To create the “noise,” dots can either be drawn in a random position in the aperture on each frame (“random position”), move to an adjacent position in a random direction in each frame (“random walk”), or have a consistent direction of motion, designated randomly at the beginning of the trial (“random direction”). Modeling our stimulus after the random-dot kinematogram from the software PsychoPy [7,8], we parameterized these different combinations of signal and noise to yield six different display options.

Additionally, dots in RDKs have a “dot lifespan,” which determines the number of frames that pass before a dot disappears and reappears somewhere else within the aperture. However, if a dot reaches the end of the aperture and its dot lifespan has not ended, then a dot can either reappear randomly in the aperture, or be reinserted from an opposite edge. We include features to customize dot lifespan length, the reinsertion rule, as well as the number of dot sets to cycle through (with each set being presented in one frame), adapting procedures from established RDK versions in MATLAB [10].

Finally, we also include links to code that integrates this RDK as a plugin with jsPsych, a library for creating and running experiments in web browsers [11]. This code can be found in the “Software location” section of this article (see below). We include parameters to control motion direction, coherence level, the total number of dots, dot size, dot color, background color, aperture shape, aperture size, the location of the aperture on the screen, the fixation cross, and how far dots move from one frame to the next. While our raw code can be implemented on any platform that uses JavaScript, incorporating our plugin with the jsPsych library may be particularly advantageous for researchers looking to use this RDK in experiments assessing reaction times;; a comparison of reaction times assessed with jsPsych and a standard software package (e.g., PsychToolbox in MATLAB), revealed that while reaction times measured by jsPsych tended to be slightly slower than PsychToolbox, response time variability was quite comparable between both software packages [12]. This indicates that response time measurements in jsPsych are sensitive enough to detect differences caused by experimental manipulations.

### Quality control

One primary concern in conducting visual psychophysical experiments on the web is that timing issues in the display could arise due to differences in internet connectivity speeds, monitor types (i.e., liquid crystal displays vs. cathode ray tubes), hardware, or web browsers used by participants. In testing our software, we have found that running the exact same code for our RDK in different browsers yields slightly different results. For example, we have identified small differences in the average frames per second when testing our RDK using Google Chrome, Firefox, and Safari on the same computer (Figure 1A). In a test of 10,000 presentations of the RDK in each browser, Chrome was the most consistent (mean: 16.632 seconds, sd: 0.364) in presenting the stimuli at the intended frame rate (16.667 seconds), followed by Firefox (mean: 17.217, sd: 0.898) and Safari (mean: 17.757, sd: 1.796).

**Figure 1.**
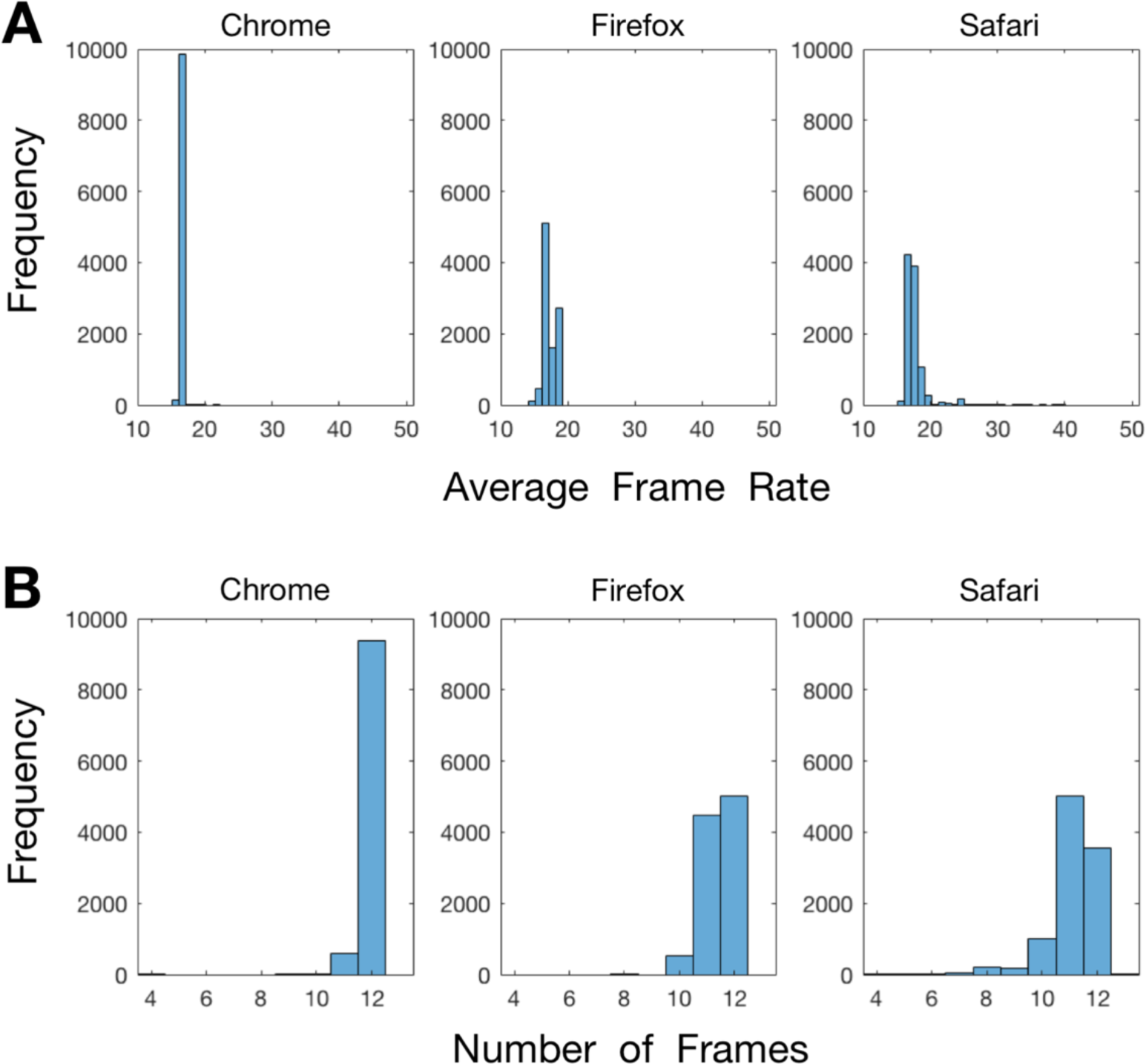
The average frame rate and number of frames shown per trial across three different web browsers. In this analysis, we presented our RDK 10,000 times in each web browser for 200ms per presentation. Each panel shows the timing differences that emerge when displaying the stimulus on the same computer in the current versions of three different web browsers: Chrome, Firefox, and Safari. Each column denotes a different browser. (A) The average frame rate for each presentation of the stimulus. In these histograms, each bin is 1ms wide and centered around the.667 mark (note that the ideal frame rate of a 60hz monitor is ~16.667ms). (B) The number of frames that were used in each 200ms stimulus presentation. The ideal number of frames is 12 using our monitor’s 60Hz refresh rate.

What is needed to ensure accuracy and precision when conducting web-based psychophysics are not only real-time measures of the average frame rate of the display, but also the number of frames that are actually presented on each trial. Our RDK records the number of frames used in each presentation, which serves as a valid index of presentation clarity and coherence (Figure 1B). We recommend that users analyze this data in one of two ways: offline, to exclude particular trials from relevant analyses, or online, to re-present trials with missed frames to ensure balanced numbers of trials across conditions. In our tests of this stimulus, Chrome had the most consistent presentation (mean: 11.937, sd: 0.258), followed by Firefox (mean: 11.448, sd: 0.594) and Safari (mean: 11.136, sd: 0.902). We did not notice visible differences between trials when 11 or 12 frames were used, but a visible stutter was apparent when the number of frames was below 10. Quality checks on other aspects of our stimulus (e.g., the locations where dots were re-drawn after disappearing, the number of dots drawn in the stimulus, whether the colors were rendered properly) demonstrated its viability, but we welcome feedback from users if issues arise.

Finally, we also recommend that users give explicit instructions to subjects to only have one browser window open when participating in experiments with this stimulus and to close other programs while it is being used, as we have noticed slight timing idiosyncrasies that emerge when other browser tabs and programs are running in the background.

## (2) Availability

### Operating system

This plugin is functional in the most recent versions of Safari, Firefox, and Chrome (i.e., the most up-to-date versions available for use in November, 2017). However, based on our tests, Chrome appears to be the most reliable browser in presenting an equal number of frames across trials for this stimulus. We welcome feedback from users if compatibility issues exist in older web browsers.

### Programming language

JavaScript/CSS/HTML

### Additional system requirements

None

### Dependencies

The raw code for the RDK posted on CodePen works without any additional frameworks or libraries. The jsPsych RDK plugin requires the jsPsych library, jQuery, and a jsPsych CSS stylesheet to work properly. These scripts and links for the jsPsych version are added in the header of the main experiment file (see the identifier of (3) under ‘Software location’ below).

### List of contributors

Sivananda Rajananda - Development & Design

Hakwan Lau - Design

Brian Odegaard - Design & Code Review

### Software location

1. Demonstration of the RDK stimulus with modifiable parameters Code repository

> **Name:** CodePen
>
> **Identifier:** https://codepen.io/vrsivananda/pen/xLORQe
>
> **Licence:** GNU General Public License version 3
>
> **Date published:** 14/09/17
>
> **Description:** This CodePen link provides users with a visual demonstration of different stimulus attributes. By changing the RDK parameters, users can see (for example) how different signal & noise rules change the appearance of the stimulus. We recommend that users change values in the “Set Parameters” section of the code to visualize different features of this stimulus.
2. jsPsych plugin for use in visual experiments Code repository

> **Name: GitHub**
>
> **Identifier:** https://github.com/vrsivananda/RDK/blob/master/jspsych-5.0.3/plugins/jspsych-RDK.js
>
> **Licence:** GNU General Public License version 3
>
> **Date published:** 14/09/17
>
> **Description:** This link leads to our RDK code that is integrated as jsPsych plugin. Please note that this jsPsych plugin is unable to work on its own, and requires use of the jsPsych framework to function properly. The user can download the raw code by clicking the “Raw” button and then copy-pasting the code, or simply right-clicking and selecting “Save as.”
3. Sample jsPsych experiment implementing the RDK stimulus Code repository

> **Name:** GitHub
>
> **Identifier:** https://github.com/vrsivananda/RDK.git
>
> **Licence:** GNU General Public License version 3
>
> **Date published:** 14/09/17
>
> **Description:** This link leads to our repository on GitHub with an example experiment that shows a functional implementation of our jsPsych-integrated RDK plugin.

### Language

JavaScript/CSS/HTML

## (3) Reuse potential

Recently, platforms have been created which make it possible to conduct psychological and perceptual experiments on the web, including the “jsPsych” JavaScript library [11] and Amazon’s mTurk website [13] for subject recruitment. RDKs are a commonly-used stimulus to evaluate motion perception thresholds in vision research;; our release of the first open access RDK for web-based experiments will be of use to any researcher interested in investigating motion perception in large numbers of subjects.

We think use of this stimulus in web-based experiments will be of particular value to researchers interested in studying populations that may be difficult to access (e.g., the elderly, individuals in various worldwide locations, individuals living far from testing sites, etc.). This RDK also facilitates fast and efficient data collection in motion perception experiments. In addition, researchers will be able to develop and extend our code under the GNU GPLv3 license to improve or modify the RDK to serve specific research purposes. We hope our creation here is of use to the greater vision science community and researchers interested in studying motion perception, and that efforts can be taken to adapt other types of visual psychophysical stimuli (Gabor patches, etc.) for web-based presentation as well.

## Competing interests

The authors declare that they have no competing interests.

